# Glutamine deprivation alters TGF-β signaling in hepatocellular carcinoma

**DOI:** 10.1101/2024.03.17.585424

**Authors:** Caroline Gélabert, Sabrina Campisano, Irene C. Golán, Nateneal T. Beyene, Carl-Henrik Heldin, Andrea Chisari, Patricia Sancho, Aristidis Moustakas, Laia Caja

**Author notes:** To whom correspondence should be addressed: Department of Medical Biochemistry and Microbiology, Science for Life Laboratory, Uppsala University, SE-751 23 Uppsala, Sweden.

## Abstract

Metabolic reprogramming is one of the hallmarks of cancer. Glutamine is one of the most important nutrients that fuels the TCA cycle and therefore takes part in the production of energy. Glutamine is used as starting metabolite for the synthesis of nucleotides, fatty acids and non-essential amino acids. Since nutrients are uptaken from the blood stream, and considering the 3-dimensional state of solid tumors, access of nutrients is highly dependent on the location of individual cells within a tumor, which results in affecting their metabolic activity. This gives rise to two disctincts cell population: the ones that have access to nutrient and the ones that are nutrient-deprived. We studied the effect of the lack of glutamine by creating glutamine-resistent hepatocellular carcinoma cell lines chosen based on their epithelial (Hep3B) or mesenchymal phenotype (SNU-499 and HLF). We found that glutamine deprivation decreased the proliferation rate, clonogenicity and stemness frequency of the three cell lines but in a greater extent of the mesenchymal cells. Transcriptomic analysis performed in HLF cells showed that glutamine deprivation decreased the activation of signaling pathways involved in cell-cell junction, cell-extracellular matrix interactions and decreased the expression of the hallmarks of epithelial-to-mesenchymal transition. We therefore investigated the role of TGFβ, a master regulator of these three processes, by transcriptomic and functional analyses in epithelial (Hep3B) and mesenchymal cells (HLF). We found that the lack of glutamine strongly impared the activation of TGFβ signaling which correlated with an altered regulation of TGFβ target genes: the expression of mesenchymal genes was no longer induced by TGFβ while the epithelial genes were more strongly induced. Functional analyses showed that glutamine deprivation abolished the invasive capacities of HCCs and decreased cell adhesion. Altogehter, our results show that glutamine metabolism is necessary to maintain a mesenchymal phenotype and to maintain an efficient TGFβ signaling in hepatocellularcarcinoma.

## Introduction

Tumor initiation and progression require the metabolic reprogramming of cancer cells in order to meet their increased bioenergetics, biosynthetic and redox demand. The metabolic reprogramming of cancer cells is today admitted as a hallmark of cancer (Hanahan & Weinberg, 2011), and was first described by Otto Warburg who showed that cancer cells increase their glucose uptake to produce energy via lactic acid fermentation instead of oxidative phosphorylation in the presence of oxygen (Warburg, 1956). In addition to glucose, glutamine is the second principal nutrient that supports survival and biosynthesis in mammalian cells (Fan *et al*, 2013). During catabolism of many amino acids, after their transamination to glutamate, the latter is converted to glutamine via glutamine synthetase (GS, alternatively known as glutamate-ammonia ligase, GLUL) and is then used as the universal transporter of amino-groups towards the liver, where the urea cycle generates the terminal catabolite urea prior to its excretion from the body (Haussinger & Schliess, 2007). Glutamine is catabolized via a process known as glutaminolysis after transport into the mitochondria and conversion into glutamate by glutaminases (GLS) (Yoo *et al*, 2020). Mitochondrial glutamate is then converted into α-ketoglutarate and free ammonia by glutamate dehydrogenase and feeds into the tricarboxylic acid (TCA) cycle in order to produce ATP (Yoo *et al*, 2020). In addition to its energetic importance, glutamine is used to synthesize other non-essential amino acids, glutathione or purine- and pyrimidine-based nucleotides (Yoo *et al*, 2020). Furthermore, glutamine is a well understood precursor to both excitatory (e.g. glutamate) and inhibitory (e.g. γ-amino-butyric acid) neurotransmitters that control neuronal activity at synapses (Albrecht *et al*, 2010).

In vivo, important nutrients, such as glucose and glutamine, are supplied by catabolism of carbohydrates and proteins respectively consumed via diet, they are synthesized in many cells in the body and circulate in the bloodstream before being uptaken by energetically active cells. In the context of solid tumors, accessibility of a given tumor cell to nutrients is modulated by its proximity to the vasculature. Cells located adjacent to the vasculature have access to nutrients and oxygen to fuel anabolic pathways that support proliferation, whereas, cells distant from the vasculature have diminished access to nutrients and oxygen (Boroughs & DeBerardinis, 2015). Such cancer cells deprived from nutrients and oxygen are usually forced to adjust their nutrient uptake and the use of metabolic pathways in order to maintain viability and malignancy.

In terms of metabolic adaptation, nutrient-deprived cells first decrease their demand for ATP in order to maintain an adequate ATP/ADP ratio (Gameiro & Struhl, 2018). If the decrease of metabolic activity is not sufficient to maintain this ratio, cells can activate certain enzymes such as adenylate kinase that converts two molecules of ADP, the precursor of ATP, into one molecule each of AMP and ATP (Klepinin *et al*, 2020). In addition to a lower metabolic activity, nutrient-deprived cells can decrease their mammalian target of rapamycin (mTOR) kinase activity in order to increase autophagy and recycle existing proteins, thus providing an intracellular glutamine supply to sustain mitochondrial function (Duran *et al*, 2012). Finally, certain Ras-transformed cancer cells counteract the lack of glutamine by the degradation of unsaturated fatty acids to support ATP production (Kamphorst *et al*, 2013). The effect of the lack of glutamine on cancer cell aggressiveness has been studied by transient glutamine deprivation followed by functional analyses. It was reported that glutamine deprivation reduces the migratory and invasive properties of ovarian cancer cells via the downregulation of matrix metalloproteases (MMPs) via the transcriptional regulator of *MMP* gene expression, ETS1 (Prasad & Roy, 2021). Cancer-associated fibroblasts (CAFs) deprived from glutamine are able to remodel the signaling complex of Akt2/TRAF6/p62 that controls polarization, in order to direct CAF mobility toward glutamine (Mestre-Farrera *et al*, 2021). In cervical carcinoma HeLa and in breast cancer cells, extracellular matrix stiffness increases microtubule glutamylation leading to increased cell invasion (Torrino *et al*, 2021). Thus, glutamine, via its direct derivative, glutamate, fuels the substrate for such glutamylation reactions.

Hepatocellular carcinoma (HCC) is the fifth most common tumor worldwide and comprises 75%-85% of cases of liver cancer (Bray *et al*, 2018). It is an inflammation-induced cancer modulated by external signals, such as the transforming growth factor beta (TGFβ). It is also reported that TGFβ plays a central role in metastasis, inflammation, fibrogenesis and immunomodulation in the HCC microenvironment (Jin *et al*, 2022). As part of its pro-tumorigenic actions, TGFβ induces the process of epithelial-to-mesenchymal transition (EMT) which correlates with an increase of the stem-related genes in different HCC cell lines (Malfettone et al 2017). This process is mediated by downstream signaling effectors, the Smad proteins and associated protein or kinases that regulate important genes that drive the EMT, such as Snail, a major EMT transcription factor, and is further mediated by crosstalk between TGFβ and other pathways or other transcription factors, such as the nuclear Liver X receptor α (LXRα) that potentiates the TGFβ effects via the regulation of Snail expression (Bellomo et al 2017). As a mechanism that sustains cancer cell malignancy, TGFβ contributes to the reprogramming of glutamine metabolism via up-regulation of the transporter Solute Carrier Family 1 Member 5 (SLC1A5), responsible for the transport of glutamine inside the cell and later inside the mitochondria, as well as the up-regulation of the enzyme GLS1 that converts glutamine into glutamate to enhance biosynthetic use of TCA metabolites (Soukupova et al 2017). In normal liver, the regulation of glutamine metabolism is essential for the maintenance of bicarbonate and the recycling of ammonia by glutamine synthetase in order to protect the liver from ammonia toxicity (Haussinger & Schliess, 2007). In this paper, we first investigated the response of HCC cells after adaptation to glutamine starvation. Then, we studied the importance of glutamine deprivation on TGFβ signaling.

## MATERIALS AND METHODS

### Cell culture

Human hepatocellular carcinoma cell lines (HLF, SNU-499, Hep3B) were cultured in Dulbeccós modified Eaglés medium without glucose (DMEM; Sigma-Aldrich AB, Stockholm, Sweden) supplemented with 10% fetal bovine serum (FBS; Biowest, Almeco A/S, Esbjerg, Denmark). The control cells were supplemented with 2 mM of glutamine and the respective concentration of glucose: HLF cells were supplemented with 25 mM of glucose, SNU-499 with 12 mM of glucose and Hep3B with 5.6 mM of glucose based on the standard cell culture protocol defined by previous investigators and/or the Ammerican Type Cell Collection (ATCC). The concentration of glutamine was gradually decreased to 0 mM to create glutamine-deprived cells. All cell lines were kept in humidified incubators at 37°C and 5% CO_2_.

### Cell proliferation assay

Cell proliferation was assessed using the PrestoBlue Cell Viability Reagent (ThermoFisher Scientific, Stockholm, Sweden) and the MTS reagent (G3582, Promega Biotech AB, Nacka, Sweden). Control or glutamine-deprived HLF, SNU-499 and Hep3B cells (10,000 cells/well) were seeded in multi-well 96 plates (Corning-Falcon, ThermoFisher Scientific, Stockholm, Sweden). After three days, cells were labeled with 10 µM of MTS or PrestoBlue for 1 h at 37°C and the respective absorbance (MTS: 490 nm, PrestoBlue: 590 nm) of the metabolic product generated was quantified in an Enspire multimode plate reader (PerkinElmer Sverige AB, Upplands Väsby, Sweden).

### Colony assay

Clonogenicity was assessed using crisal violet staining of 1,000 HLF, SNU-499 or Hep3B cells seeded in multi-well 6 plates (Corning-Falcon, ThermoFisher Scientific, Stockholm, Sweden) in their approporiate culture medium (control or glutamine-deprived). Cells were incubated for three weeks at 37 °C. Cells were placed on ice, rinsed with cold PBS and fixed with ice-cold MetOH for 10 min. Cells were stained with 0.5% (w/v) crystal violet diluted in 20% (v/v) MetOH for 10 min and rinsed with ddH_2_O. The numbers of colonies were counted.

### Extreme limiting dilution assay

Control or glutamine-deprived HLF, SNU-499 and Hep3B cells were seeded on Corning Costar low-attachment 96-well plates (Corning-Costar, ThermoFisher Scientific, Stockholm, Sweden) in decreasing serial dilutions (100-3 cells/well), in 6 replicates per condition and sphere formation was analyzed after 10 days of incubation at 37°C using the online ELDA analysis program (http://bioinf.wehi.edu.au/software/elda).

### Intracellular redox state analysis

Control or glutamine-deprived HLF, SNU-499 and Hep3B cells were seeded in multi-well 12 plates (Corning-Falcon, ThermoFisher Scientific, Stockholm, Sweden). Three days later, cells were incubated with 2.5 µM DCFH-DA (2’,7’-dichloro-dihydro-fluorescein-diacetate; excitation at 495 nm, emission at 520 nm; Life Technologies, Europe BV, Stockholm, Sweden) in Hank’s buffered saline solution without phenol red for 30 min at 37°C. Intracellular esterases convert DCFH-DA to 2’,7’-dichloro-dihydro-fluorescein (DCFH) that in turn is converted into 2’,7’-dichloro-fluorescein (DCF) when oxidized by H_2_O_2_. Cells were lysed in 25 mM Hepes pH 7.5, 60 mM NaCl, 1.5 mM MgCl_2_, 0.2 mM EDTA, 1% Triton-X-100 for 10 min at room temperature followed by fluorimetry quantification using an Enspire multimode plate reader. The DCF quantification was normalized on the corresponding protein amount that was quantified using the BCAprotein assay kit (ThermoFisher Scientific, Stockholm, Sweden).

### Immunoblotting

Total proteins from HLF, SNU-499 or Hep3B cells were exracted after treatment or not with TGFβ1 (5 ng/ml) for 1 or 48 h in lysis buffer (20 mM Tris-HCl, pH 8.0, 1% nonidet-P40, 150 mM NaCl, 2 mM EDTA complemented with phosphatase ihibitor (Roche Diagnostics Scandinavia AB, Bromma, Sweden) and complete protease inhibitor mixture (Roche Diagnostics Scandinavia AB, Bromma, Sweden). Proteins were then quantified and subjected to sodium dodecyl sulphate-polyacrylamide gel electrophoresis. The resolved proteins were transferred to nitrocellulose using a wet transfer unit (Bio-Rad Laboratories AB, Solna, Sweden). The efficiency of immunoblotting and equal loading of proteins was verified by staining of the nitrocellulose membrane with 0.1% (w/v) Ponceau S in acetic acid. Upon incubation with primary antibodies and horseradish peroxidase-conjugated secondary antibodies, enhanced chemiluminescence assays were performed using the Millipore kit (Merck-Millipore, Stockholm, Sweden). The antibodies used are listed in the Supplementary Table S1. Quantification of protein signals by densitometry was performed using National Institutes of Health ImageJ software. Band densities for protein of interest were normalized to that of the band for β-actin or α-tubulin in the same sample.

### Real-time RT-PCR

The total RNA from HLF, SNU-499 and Hep3B cells was extracted using the NucleoSpin RNA kit (Macherey-Nagel, AH Diagnostics, Solna, Sweden) according to the protocol of the manufacturers. cDNA was synthesized using the iScript cDNA synthesis kit (Bio-Rad Laboratories AB, Solna, Sweden). Real-time PCR was done in triplicate using the qPCRBIO SyGreen 2×Master Mix (PCR Biosystems, London, UK) and gene-specific primers listed in the Supplemetary Table S2. Gene expression levels were determined by the comparative C_t_ method and using *HPRT1* (hypoxanthine phospho-ribosyl transferase 1) as reference gene. Normalized mRNA expression levels are plotted in bar graphs that represent average values from triplicate determinations with standard deviations. Each independent experiment was repeated at least three times.

### Cell invasion assay

The invasion assay was designed using transwell plates with 6.5 mm diameter and 8 μm pore filters (351152, Corning Costar, NY, USA or 140629, Thermo Fischer Scientific, Uppsala, Sweden). Inserts were coated with 100 μg/ml collagen (5005, Advanced BioMatrix Inc., San Diego, CA, USA) and incubated overnight at 37 °C. The HLF or Hep3B cells (5×10^4^) were seeded in the upper chamber coated with collagen. For all cell lines, cells were seeded in the upper chamber in serum-free DMEM complemented with the appropriate nutrient concentration and treated or not with TGFβ1 (5 ng/ml) during the migration time, and DMEM/6% FBS was placed in the lower chamber as chemoattractant to invading cells. Following 15 h, inserts were fixed with ice-cold MetOH. Nuclei were stained with 4′,6-diamidino-2-phenylindole (DAPI) 1:1,000 (Sigma-Aldrich Sweden AB, Stockholm, Sweden). Invasion of the cells was measured by counting the nuclei of the cells that migrated through the filter pores towards the chemoattractant (6% FBS). For quantification, 10 pictures of each insert were taken at 20× magnification, and nuclei were counted using the ImageJ software (National Institutes of Health, Bethesda, MD, USA). Results are expressed in number of cells per picture.

### Extracellular matrix-mediated cell adhesion

Cells (15,000 per well) were seeded in 96-well plates coated with the extracelular matrix (ECM) proteins collagen I (1.7 mg/ml, PureCol, Advanced BioMatrix, Inc., San Diego, CA, USA) or without coating for 60 min. Pluronic F-127 (10 mg/ml, Sigma-Aldrich AB, Stockholm, Sweden) was used to block any uncoated surface of the coated wells prior to cell seeding. Adherent cells were stained with crystal violet diluted in 20% (v/v) MetOH and dried overnight at 37°C. Crystal violet was re-diluted in pure MetOH at room temperature for 30 min and quantified at absorbance 570 nm in an Enspire multimode plate reader (PerkinElmer Sverige AB, Upplands Väsby, Sweden).

### Senescence assay

Senescent cells were detected using a senescence β-Galactosidase staining kit (9860; Cell Signaling Technology/BioNordika Sweden, Solna, Sweden). Cells (10,000 per well) were seeded in a multi-well 12 plate in control or glutamine-deprived medium and treated or not with TGFβ1 (5ng/mL) and stainined three days after seeding according to the manufactureŕs instructions. β-Galactosidase staining was observed under phase-contrast microscopy and cells were photographed in a phase contrast microscope (Nikon-Eclipse Ti-U, Nikon Europe B.V., Amstelveen, The Netherlands), equipped with a CCD camera (Andor multi pixel sCMOS camera, Oxford Instruments, Abingdon, UK).

### RNA-seq

1 microgram total RNA extracted using ReliaPrep™ RNA Cell Miniprep System (Promega, Madison, WI, USA), were checked using the Agilent-2100 Bioanalyzer System and subjected to library preparation utilizing the TruSeq stranded total RNA Gold library preparation kit with RiboZero Gold treatment and unique dual indexes according to the manufacturer’s instructions (Illumina Inc., San Diego, CA, USA). The paired-end reads of 150 bp were generated in an S4 flowcell with v1.5 sequencing chemistry on a NovaSeq-6 000 platform (Illumina Inc.). Read quality was checked by FastQC tool v0.11.9, reads were aligned against the reference human genome (GRCh38) using STAR aligner v2.7.2b with two-pass mode (Dobin *et al*, 2013). After checking alignment quality by SAMStat tool v1.5.1 (Lassmann *et al*, 2011), high-quality (Q = 30) aligned reads were annotated and quantified against the gencode comprehensive gene annotation release 38 (GRCh38.p13) using featureCounts of subread package v2.0.0 (Liao *et al*, 2014). Differential gene expression analysis was performed by EdgeR Bioconductor package in R (Chen *et al*, 2016) thus calculating normalized counts per million reads. We considered cut-off criteria of log_2_ fold-change ±1 and false discovery rate (FDR) < 0.05 for differential expression. Functional enrichment of differentially expressed genes was based on the R package clusterProfiler (http://www.bioconductor.org/) (Yu *et al*, 2012).

### Statistical analysis

All figure legends and each detailed method present the biological and technical replicates and the assessment of statistical significance. The statistical method used was based on sample content and variation within each data-set; similar variance existed between the compared group populations.

## RESULTS

### Glutamine deprivation affects the morphology and glutamine metabolism of mesenchymal HCC

In order to study the importance of glutamine metabolism in HCC cell aggressiveness, we created stable, glutamine-deprived HCC cell lines. Three cell lines were selected according to their mesenchymal or epithelial features, in order to capture the inherent heterogeneity of HCC tumor types: the HLF that exhibits the most mesenchymal phenotype, the SNU-499 with intermediate phenotype and the more epithelial Hep3B cell line (Caruso *et al*, 2019, Coulouarn *et al*, 2008). These cell lines were grown in DMEM medium supplemented with 2 mM glutamine. Then, the concentration of glutamine was decreased gradually throughout the cell passages to reach a concentration of 0 mM of glutamine (Fig. 1A). The morphology of the HLF glutamine-deprived cells was very different from the control, which are elongated and individualized, while the glutamine-deprived cells were rounded up and developed into clusters (Fig. 1B). Similarly to the HLF but to a lesser extent, the mensechymal morphology of the SNU-499 control was altered by the lack of glutamine (Fig.1C). In contrast, the epithelial Hep3B did not show a significant change in morphology (Fig. 1D).

**Figure 1.**
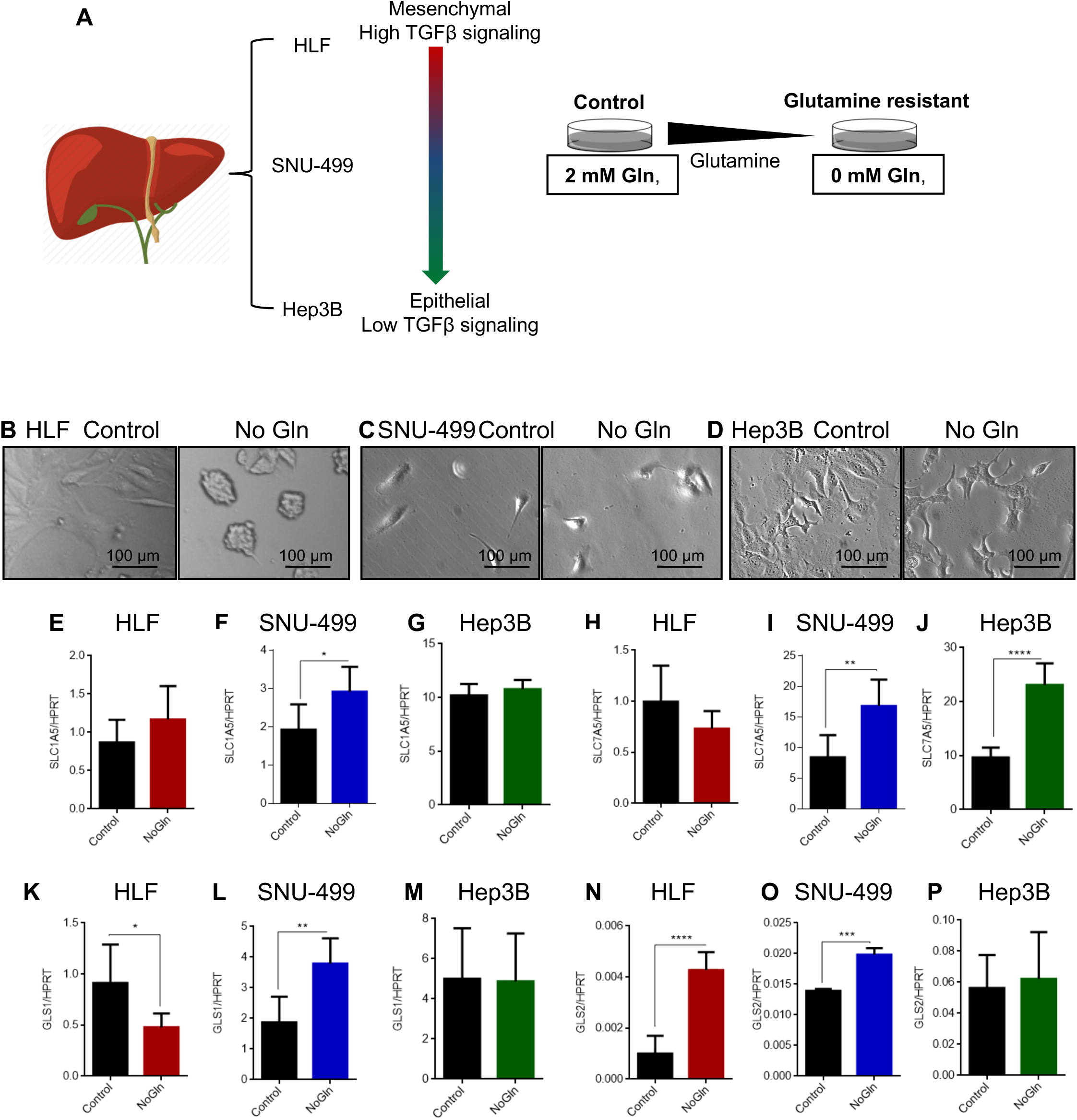
Experimental approach and general characterisation of glutamine-deprived hepatocellular carcinoma cell lines. A. Illustration of the experimental approach to study the effect of glutamine deprivation in hepatocellular carcinoma cell lines. B-D. Phase-contrast images showing the HLF (B), SNU-499 (C) and Hep3B (D) cell lines under control or glutamine-deprived (no Gln) culture conditions. Magnification bars (100 μm). E-G. RT-qPCR for determination of the basal expression of the transporter SLC1A5 in the indicated cell line, cells under control or glutamine-deprived (no Gln) culture conditions. H-J. RT-qPCR for determination of the basal expression of the transporter SLC7A5 in the indicated cell line, cells under control or glutamine-deprived (no Gln) culture conditions. K-M. RT-qPCR for determination of the basal expression of the enzyme GLS1 in the indicated cell lines under control or glutamine-deprived (no Gln) culture conditions. N-P. RT-qPCR for determination of the basal expression of the enzyme GLS2 in the indicated cell lines under control or glutamine-deprived (no Gln) culture conditions. RT-qPCR graphs present data from (n=3) biological repeats as means with SEM and associated significance as: * p<0.05, ** p<0.01, *** p<0.001, **** p<0.0001.

Glutamine uptake by mammalian cells is mediated by various transporters on the cell surface and upon uptake inside the cells, glutamine is converted to glutamate, which then enters the TCA cycle (Yoo *et al*, 2020). The transporter SLC1A5 (ASCT2) that regulates the flux of glutamine into cells was up-regulated in HLF and SNU-499 cells adapted to the absence of glutamine (Fig. 1E, F), but not in the equivalent Hep3B cells (Fig. 1G). In contrast, the transporter SLC7A5 that regulates the entry of all amino acids was equally expressed in both parental and glutamine-deprived HLF cells (Fig. 1H), but up-regulated in glutamine-deprived SNU-499 and Hep3B cells (Fig. 1I, J). We then examined the expression of the intracellular enzymes responsible for the conversion of glutamine to glutamate, GLS1 being expressed in all cell types and GLS2 being restricted to liver cells (Yoo *et al*, 2020). GLS1 and GLS2 showed a different expression profile among the three cell lines. The most abundant GLS1 enzyme was slightly down-regulated in the HLF glutamine-deprived cells (Fig. 1K), up-regulated in the SNU-499 glutamine-deprived cells (Fig. 1L) and not affected in the glutamine-deprived Hep3B cells (Fig. 1M). In contrast, the liver-specific GLS2 enzyme was up-regulated in the two mesenchymal cell lines HLF and SNU-499 (Fig. 1N,O), but remained unchanged in the epithelial cell line Hep3B (Fig. 1P). The different adaptation to glutamine deprivation among these three cell lines showed a greater dependency of the mesenchymal cells to this amino acid compared to the epithelial cells, the former apparently trying to compensate the lack of external glutamine supply by adjusting the expression of GLS enzymes.

### Glutamine deprivation diminishes cell viability, clonogenicity, stemness frequency and ROS production

By converting to glutamate, glutamine becomes a key nutrient for ATP production and therefore important for cell proliferation and viability. Cell proliferation and viability were assessed with two independent but similar methods to quantify proliferation, the PrestoBlue reagent and the MTS reagent, both reduced by viable cells into metabolites that can be quantified spectrophotometrically. Both methods showed significantly lower numbers of viable, proliferating cells in the three glutamine-deprived cell lines (Fig.2A-F). We also analyzed a key regulator of the cell cycle, cyclin E, in these cells. HLF glutamine-deprived cells consistently showed a decrease in Cyclin E expression (Supplementary Fig. 1A). The level of Cyclin E remained unchanged in Hep3B glutamine-deprived cells (Supplementary Fig. 1B), which can be explained by an already low proliferative rate in the control condition, or by regulation at the level of cyclin-dependent kinase function. Although cell proliferation and viability was diminished in glutamine-deprived cells, they did not show clear signs of senescence, measured as β-galactosidase-positive cells (Supplementary Fig. 1 C, D), or apoptosis, shown by the absence of cleaved caspase3 or PARP1 (Supplementary Fig. 1 A, B), as the levels of these proteins were similar both in control and glutamine-deprived cells.

**Figure 2.**
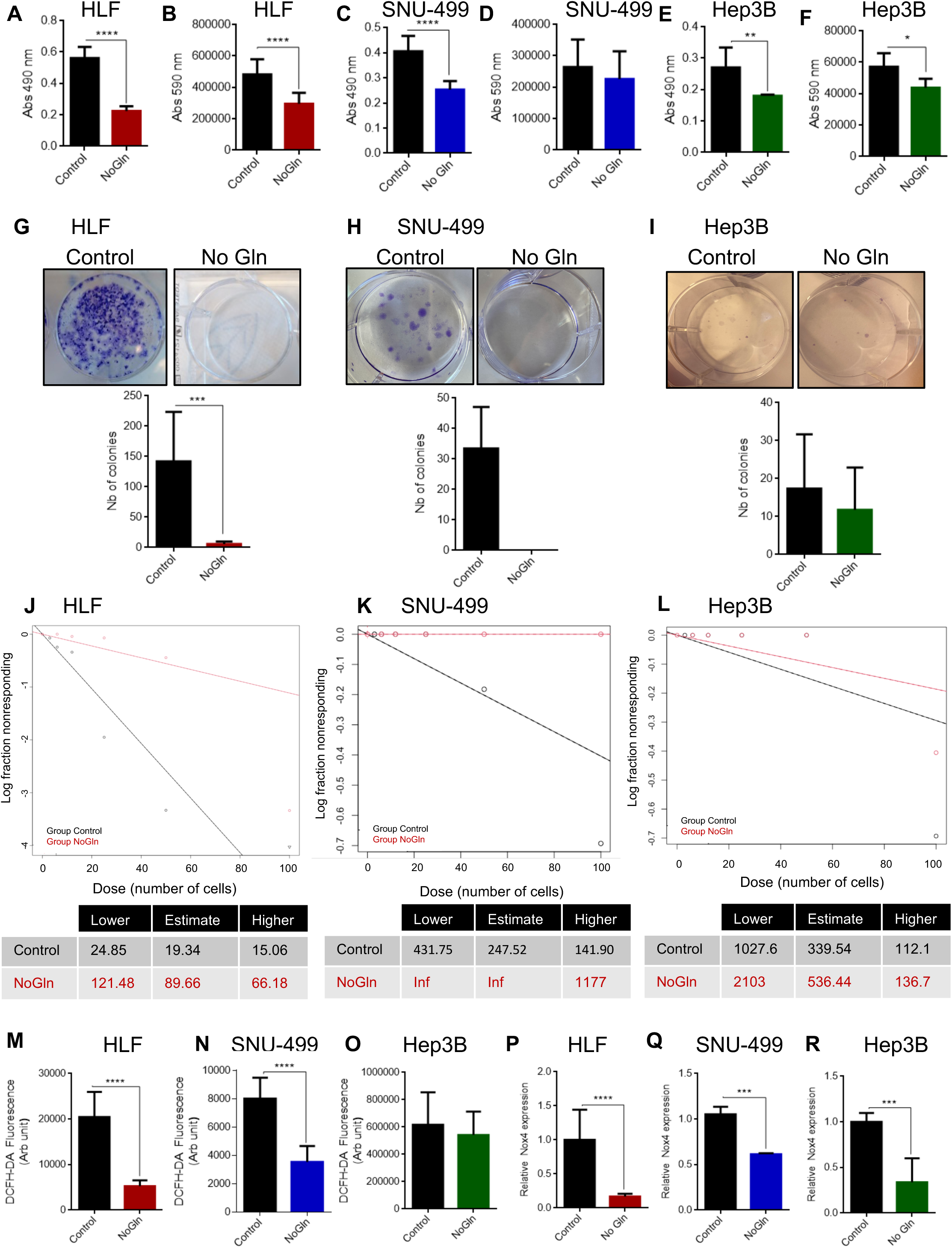
Glutamine deprivation decreases cell viability, colony formation capacities, self-renewal frequency and ROS production in HCC. A-F. Cell viability measured in the indicated cell lines under control or glutamine-deprived (no Gln) culture conditions by PrestoBlue assay (Abs 490 nm. A, C, E) or MTS assay (Abs 590 nm, B, D, F) 72 h after seeding. G-I. Colony formation assay determined by crystal violet staining 3 weeks after seeding in the indicated cell line under control or glutamine-deprived (no Gln) culture conditions. J-L. Self-renewal frequency determined by ELDA. Graphs show the median values in control or glutamine-deprived cells (Control: black curves; NoGln: red curves) in the indicated cell lines. High x-axis intercept corresponds to low number of spheres. The table shows the self-renewal frequency (1 stem cell/x cells) or Inf: too low frequency. For HLF and SNU-499: n = 3 with six replicates, and for Hep3B: n = 1 with six replicates. M-O. Levels of intracellular ROS measured by DCFH-DA in control or glutamine-deprived cells in the indicated cell lines three days after seeding. P-R. RT-qPCR for determination of *NOX4* mRNA expression in control or glutamine-deprived cells in the indicated cell lines. Each graph represents data from (n=3) biological replicates as means with SEM and associated significance as: * p<0.05, ** p<0.01, *** p<0.001, **** p<0.0001.

The capacity of single HLF, SNU-499 and Hep3B cells to produce a colony of descendent cells was assessed by clonogenic assays on an adherent substrate. The HLF and SNU-499 control cells exhibited a high clonogenic capacity (150 and 32 colonies per well respectively) but no colonies were formed by the glutamine-deprived cells (Fig. 2G, H). Unlike the HLF and SNU-499 cells, the Hep3B cells were form very few colonies, either in the control or in the absence of glutamine (Fig. 2I). The colony forming capacity of the cells reflects their proliferative rate but also their self-renewal potency. The self-renewal frequency of the cells was assessed by extreme dilution assay on low-attachment substrate. The lack of glutamine caused a very reproducible and significant decrease of sphere formation in the mesenchymal cells HLF (Fig. 2J, Supplementary fig. 2A) and SNU-499 (Fig. 2K, Supplementary fig. 2B), from 1/19 HLF cells in the control condition to 1/89 HLF cells in the glutamine-deprived condition, and 1/146 SNU-499 cells in the control condition to 1/609 SNU-499 cells in the glutamine-deprived condition. The Hep3B cells were unable to form spheres either in control or glutamine-deprived conditions (Fig. 2L, Supplementary fig. 2C) which can be explained by their strong epithelial phenotype associated with a more differentiated state.

Cell metabolism involves the mitochondrial TCA cycle and the electron transport pathway to produce the energy needed for the cells to proliferate. The mitochondrial respiratory chain produces ATP through a transmembrane electrochemical gradient. The final electron recipient of the mitochondrial chain is molecular oxygen (O_2_), which can be incompletely reduced giving rise to different forms of oxygen derivatives. Oxygen-derived free radicals are called reactive oxygen species (ROS). Cancer cells contain elevated quantities of ROS due to their high proliferative rate and cancer cell ROS production has been associated with various cellular responses (Mazat *et al*, 2020). In agreement with this statement, the glutamine-deprived HLF and SNU-499 cells, which exhibit a lower proliferative rate, also produced less intracellular ROS than the respective control cells (Fig. 2M, N). However, the Hep3B cells that grew very slowly dit not show any difference in their ROS production (Fig. 2O), despite the reduction of proliferation rate in the glutamine-deprived cells (Fig. 2E, F). In addition to the respiratory chain contribution to ROS synthesis, the elevation of ROS levels can be contributed to the NADPH-oxidase family (Nox) of proteins. Cancer stem cells are known to have high basal levels of ROS, and among other Nox members, NOX4, which is constitutively active in cells, is known to regulate the expression of cancer stem cell genes in glioblastoma (Garcia-Gomez *et al*, 2022) and promote oncogenesis in pancreatic cancer (Ju *et al*, 2017). On the other hand, NOX4 has been reported to act as a tumour suppressor or oncogene depending on the tumour type. In HCC, NOX4 has been described to prevent cancer progression (Crosas-Molist *et al*, 2014, Penuelas-Haro *et al*, 2022). For these reasons we analyzed Nox4 expression, and observed reduced *Nox4* mRNA levels in all three cell lines upon glutamine deprivation (Fig. 2P-R). The decrease of *Nox4* expression in the glutamine-deprived cells therefore correlates with the decrease of self-renewal frequency observed in HLF and SNU-449 cells (Fig. 2J-L). The lack of change in ROS levels in Hep3B cells (Fig. 2O) also suggests that the observed reduction in *Nox4* mRNA expression in the same cells upon glutamine deprivation (Fig. 2R) may have alternative functional roles.

Altogether, these results show a decrease of cell proliferation in the three cell lines, and a decrease of clonogenicity, self-renewal frequency and intracellular ROS production in the two mesenchymal cell lines HLF and SNU-499 but not in the epithelial cell line Hep3B, consistently with Fig. 1 that suggested an increased dependence of the mesenchymal cells on glutamine metabolism.

### Glutamine deprivation alters gene expression and TGFβ response

In order to identify differentially expressed genes in control cells compared to glutamine-deprived cells, RNA-seq analysis was performed in the HLF cell line that showed the most dramatic phenotypic changes after glutamine starvation (Fig. 1, 2). By comparing the control cells with the glutamine-deprived cells, we found differential expression (both enhanced or decreased) of several genes, indicating that glutamine deprivation affects widespread gene expression (Fig. 3A, B).

**Figure 3.**
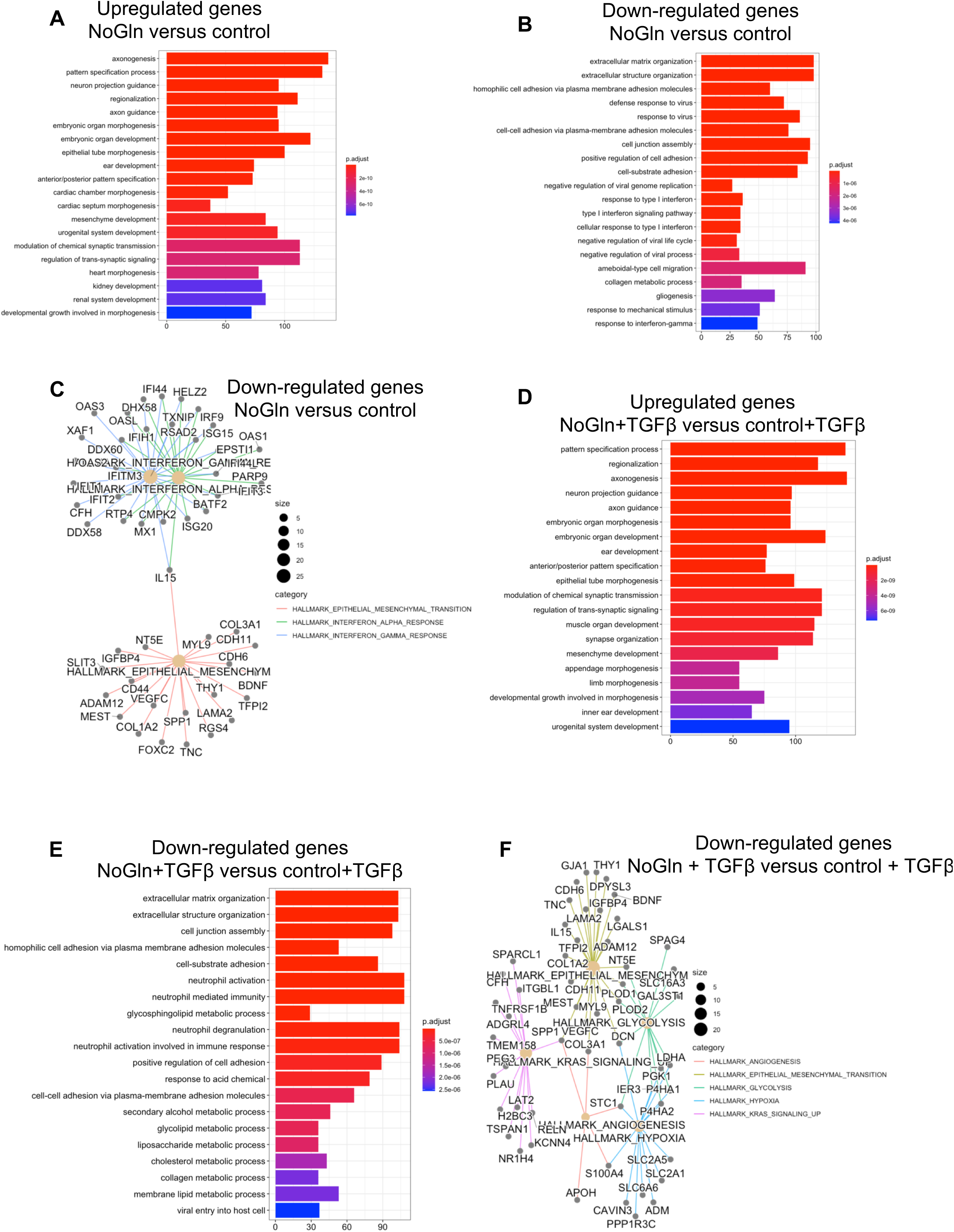
Transcriptomic analysis of glutamine-deprived HLF cells. A, B. Gene ontology (GO) analysis of the biological processes of the up- (A) or down- (B) regulated genes between control versus glutamine-deprived cells. Adjusted p-value is color-coded. C. Hallmark pathway analysis among down-regulated genes due to the lack of glutamine. D, E. GO analysis of the biological processes of the up- (D) or down- (E) regulated genes after 24 h of TGFβ treatment between control versus glutamine-deprived cells. Adjusted p-value is color-coded. F. Hallmark analysis among down-regulated genes due to the lack of glutamine after TGFβ treatment for 24 h.

More specifically, glutamine deprivation caused up-regulation of 3,895 genes and down-regulation of 3,573 genes (Supplementary Fig. 3A). Gene ontology analysis revealed that adaptation to glutamine starvation leads to the upregulation of genes related to pathways of cellular morphogenesis, including epithelial and mesenchymal morphogenesis, heart, kidney and neuronal morphogenesis (Fig. 3A). These data suggested broadly that glutamine deprivation favored a certain plasticity change in the HCC cells manifested by expression of genes involved in diverse cellular lineages or fates. In contrast, the down-regulated genes enriched for molecular signatures mostly related to ECM organization and structure, cell adhesion and responses to interferon signaling (Fig. 3B). In addition, the hallmarks gene set database found the hallmarks of EMT, a process that involves a modification of the ECM organization and cell adhesion, as significantly down-regulated in the glutamine-deprived cells (Fig. 3C).

The reorganization of the ECM is very important for cancer cells to create their appropriate microenvironment (Friedl & Alexander, 2011). Furthermore, in order to migrate and invade, cancer cells need to modify their cell-cell adhesion and cell-ECM interactions (Friedl & Alexander, 2011). Such changes depend on specific signals from the cancer cell microenvironment, including the TGFβ pathway (Tsubakihara & Moustakas, 2018). It is known that paracrine and autocrine TGFβ signals are required for HCC initiation and progression via the regulation of tumor proliferation, EMT and tumor microenvironment (Caja *et al*, 2018). We therefore performed RNA-seq analysis in HLF control or glutamine-deprived cells after TGFβ stimulation for 24 h. Regarding the effect of TGFβ in the control condition, we observed 296 up-regulated genes and 640 down-regulated genes (Supplementary Fig. 3C) but only 59 up-regulated genes and 48 down-regulated genes after TGFβ treatment in the glutamine-deprived condition (Supplementary Fig. 3D). These results show that the lack of glutamine alters the power of TGFβ in regulating gene expression, and it is therefore possible that TGFβ signaling requires the presence of glutamine. We found similar function and number of differentially expressed genes when comparing glutamine-deprived cells treated with TGFβ to control cells treated with TGFβ (Fig. 3D, Supplementary Fig. 3B) to the ones obtained when we compared glutamine-deprived cells to control cells (Fig 3A). The up-regulated genes were related to epithelial, mesenchymal, muscle, ear, appendage and neuronal morphogenesis (Fig. 3D), while the TGFβ-induced genes related to ECM organization and cell-cell junctions scored as enriched in the less induced gene group in glutamine-deprived HLF cells (Fig. 3E). Hallmarks gene set database analysis revealed a down-regulation of the hallmarks of EMT, angiogenesis, hypoxia and K-Ras signaling in the glutamine-deprived cells after TGFβ stimulation compared to controls cells treated with TGFβ (Fig. 3F).

### TGFβ signaling and cell invasion are modulated by glutamine availability

TGFβ signaling has a very pleiotropic role in the cells and implicates the activation of various pathways (Moustakas & Heldin, 2009, Tzavlaki & Moustakas, 2020). Consistently with the RNA-seq and pathway enrichment analysis that revealed an alteration of physiological responses regulated by TGFβ such as the regulation of cell-cell junction and EMT both processes regulated by TGFβ, the expression of TGFβ related genes was found to be differentially expressed in control versus glutamine-deprived cells (Fig. 4A), glutamine starvation lead to decrease expression of SMAD2, SMAD3 and TGFBR2, while increasing the expression of TGFBR1, SMAD6 among other genes; these changes in expression of TGFβ related genes could explain why TGFβ could not regulate the expression of as many genes in the glutamine-deprived cells compared to control cells (Suppl Fig 3C-D). The effect of glutamine deprivation on TGFβ signaling was studied in the two most different cell lines, the most mesenchymal HLF and the most epithelial Hep3B. The control HLF and Hep3B cells or corresponding glutamine-deprived cells were treated with TGFβ for one hour, an optimal time point to observe the activation and phosphorylation of downstream signaling. The activation of the Smad, ERK and Akt pathways was detected by immunoblot of their total and phosphorylated forms (Fig. 4B, C). In the HLF and Hep3B cells, we observed a clear activation of the Smad pathway by TGFβ in the control condition but not in the absence of glutamine (Fig. 4B, C). In both cell lines, the expression of total Smad2 was reduced in the glutamine-deprived cells both at the protein (Fig. 4B, C) and the mRNA levels (Fig. 4D, E), as well as the Smad3 and Smad4 mRNA levels in HLF cells. We also observed a strong activation of the ERK pathway at the basal level and weak further enhancement after TGFβ stimulation, but such activation completely disappeared in the absence of glutamine in the HLF cells (Fig. 4B) and was strongly decreased in the Hep3B cells (Fig. 4C). The activation of the Akt pathway seems to be different among the two cell lines; this pathway was weakly activated in the control HLF cells and strongly induced in the absence of glutamine (Fig. 4B), while it was strongly active in the control Hep3B cells and reduced in the absence of glutamine (Fig. 4C).

**Figure 4.**
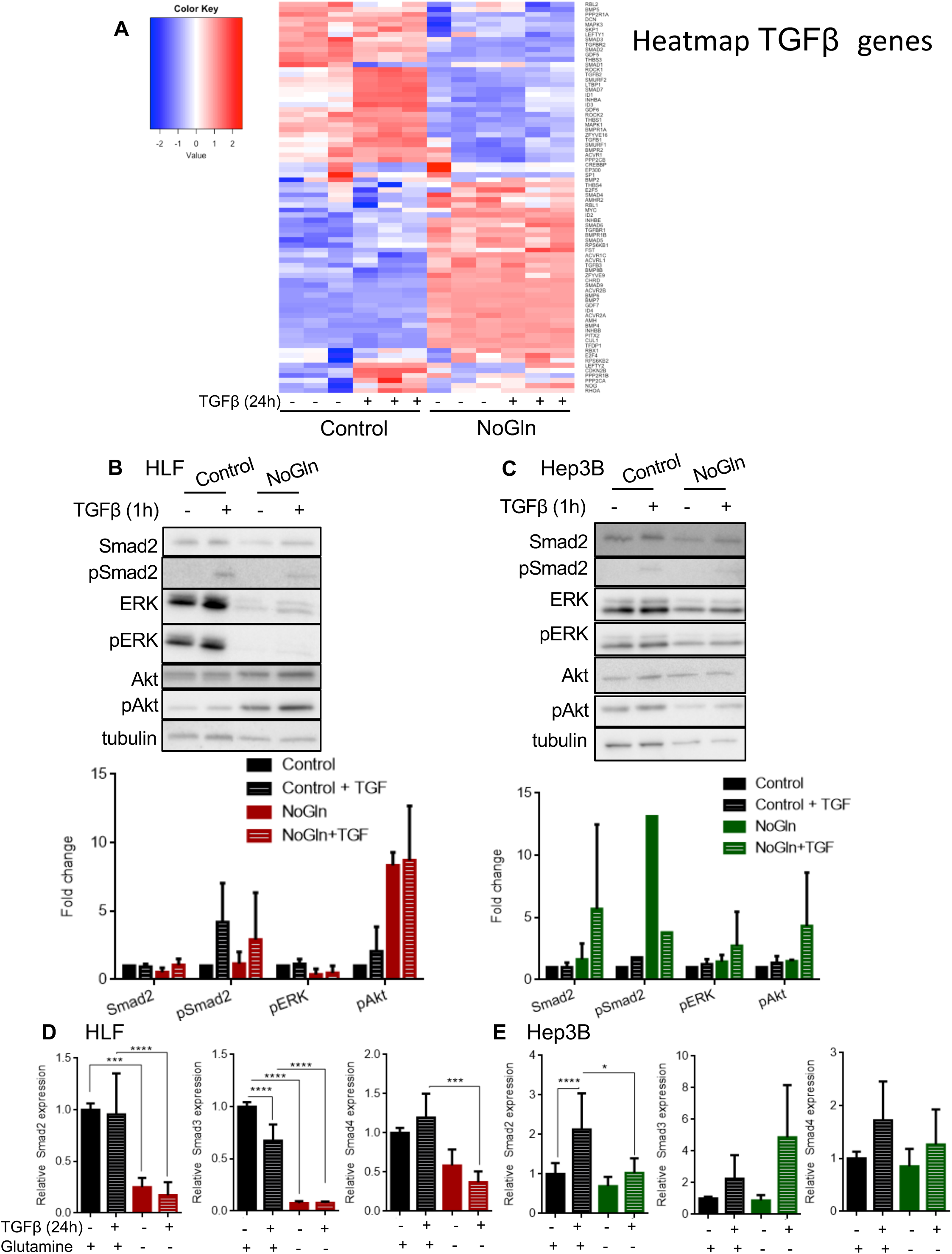
Glutamine deprivation alters TGFβ signaling in HCC. A. Heat-map of the KEGG TGFβ-related genes expressed in HLF cells control and glutamine-deprived cells treated or not with TGFβ for 24 h. B, C. Immunoblot and quantification for the indicated proteins (Smad2, pSmad2, ERK, pERK, Akt, pAkt, tubulin) in HLF (B) and Hep3B cells (C) under control or glutamine-deprived (no Gln) culture conditions after TGFβ treatment for 1 h. D, E. RT-qPCR for determination of *Smad2*, *Smad3* and *Smad4* mRNA levels in HLF (D) or Hep3B cells (E) under control or glutamine-deprived (no Gln) culture conditions. Each graph represents data from (n=3) biological replicates as means with SEM and associated significance as: * p<0.05, ** p<0.01, *** p<0.001, **** p<0.0001.

The modulation of TGFβ signaling by the lack of glutamine prompted us to investigate the effect of glutamine starvation on specific genes regulated by the TGFβ pathway. Although the morphology of the HLF and Hep3B cells were not affected by TGFβ stimulation for 48 h (Fig. 5A, B), the EMT-related genes regulated by TGFβ showed significant differences in control versus glutamine-deprived cells and among the two tested cell lines as already indicated by the RNASeq analysis (Fig 3). As expected, the mesenchymal HLF cell line grown in the presence of glutamine expressed high levels of fibronectin, vimentin and plasminogen activator inhibitor (PAI) 1 (Fig. 5C). Their level of expression being already high in the absence of TGFβ, expression levels of these proteins were not further enhanced by TGFβ, whereas the epithelial marker E-cadherin was not expressed in the control HLF cells (Fig. 5C). The expression of the EMT-related proteins was reversed in the absence of glutamine; the expression of E-cadherin was strongly induced, even in the absence of TGFβ, and the mesenchymal proteins fibronectin, vimentin and PAI1 were strongly reduced (Fig. 5C). On the other hand, the epithelial cell line Hep3B expressed E-cadherin but low levels of the mesenchymal proteins fibronectin, vimentin and PAI1 in the control condition (Fig. 5D). After TGFβ treatment for 48 h, the level of expression of E-cadherin was reduced and that of the mesenchymal proteins was induced, as expected (Fig. 5D). In the absence of glutamine, the Hep3B cells expressed higher levels of E-cadherin and the TGFβ-induced mesenchymal proteins were weaker than the control (Fig. 5D). These results were confirmed by immunofluorescence that showed a lower expression of fibronectin, vimentin and vinculin by TGFβ and an increased expression of E-cadherin in HLF cells deprived from glutamine (Supplementary Fig. 4). Consistently with the protein level, the mRNA levels of *E-cadherin* were enhanced and the mRNA levels of the mesenchymal genes such as *N-cadherin*, *Snail* and *fibronectin* were reduced in the glutamine-deprived HLF cells (Fig. 5E). However, the expression of E-cadherin at the protein and mRNA levels did not correlate in Hep3B cells, which possibly suggests differential regulation of the stability of the protein in control versus glutamine-deprived cells (Fig. 5F). The mesenchymal markers *N-cadherin* and *fibronectin*, but not *Snail*, were reduced in the absence of glutamine expression (Fig. 5F). Altogether, these results show that the lack of glutamine affects TGFβ signaling activation and pushes the cells to become more epithelial, suggesting that glutamine metabolism is needed to maintain a mesenchymal phenotype.

**Figure 5.**
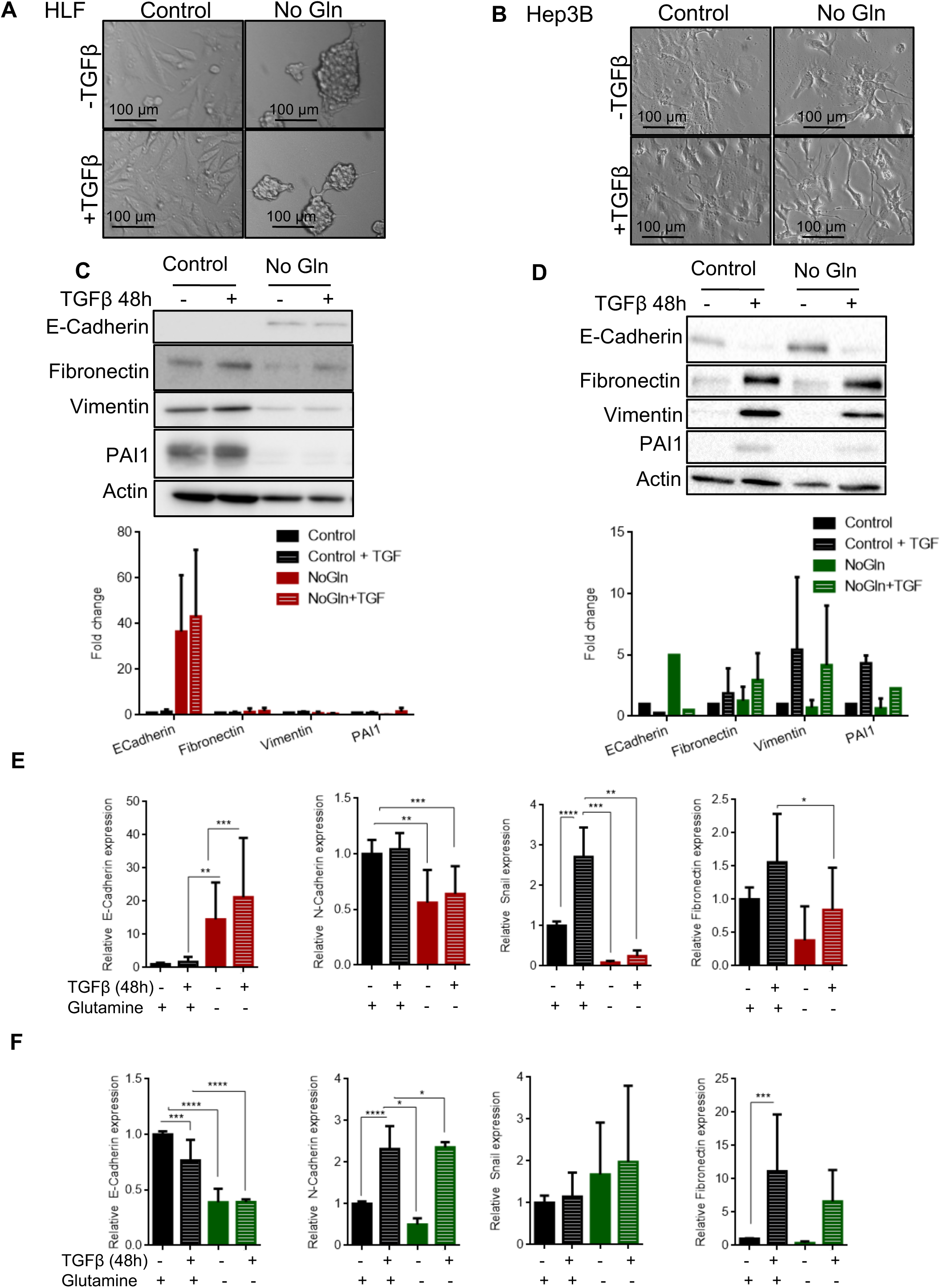
Glutamine deprivation alters EMT-related genes induced by TGFβ in HCC. A, B. Phase-contrast images showing the HLF (A) and Hep3B (B) cell lines under control or glutamine-deprived (no Gln) culture conditions and treated or not with TGFβ for 24 h. Magnification bars (100 μm). C, D. Immunoblot and quantification for the indicated proteins (E-cadherin, fibronectin, vimentin, PAI1, β-actin) in HLF (C) and Hep3B cells (D) under control or glutamine-deprived (no Gln) culture conditions after TGFβ treatment for 48 h. E, F. RT-qPCR for determination of *E-cadherin*, *fibronectin*, *vimentin*, *PAI1* mRNA levels in HLF (E) and Hep3B cells (F) under control or glutamine-deprived (no Gln) culture conditions after TGFβ treatment for 48 h. Each graph represents data from (n=3) biological replicates as means with SEM and associated significance as: * p<0.05, ** p<0.01, *** p<0.001, **** p<0.0001.

The comparison of RNA-seq data showed a differential expression of genes involved in EMT (Figure 6A) and focal adhesion assembly (Fig. 6B). These results were corroborated by functional assays. The invasive capacities of HLF and Hep3B cells were investigated using collagen-based transwell assays. Consistent with the expression of the EMT genes, the invasiveness capacity of the HLF cells was not significantly enhanced by TGFβ stimulation (Fig. 6C), unlike Hep3B cells that strongly responded to TGFβ (Fig. 6D). However, the absence of glutamine completely abolished the invasive capacity of both cell lines, in the absence or presence of TGFβ (Fig. 6C, D). The loss of the invasive capacity of the glutamine-deprived cells corresponded to a decrease in cell adhesion by both cell lines (Fig. 6E, G). At the molecular level, the activated focal adhesion kinase (FAK) visualized as phosphorylated FAK (pFAK) and the activated Src visualized as phosphorylated Src (pSrc), activations known to be associated with metastasis and poor prognosis (Wendt & Schiemann, 2009), were no longer activated in the HLF glutamine-deprived cells (Fig. 6F, H). It is therefore clear, that glutamine is a necessary factor for propagation of the mesenchymal HCC cell fate and the associated cell adhesion and motility.

**Figure 6.**
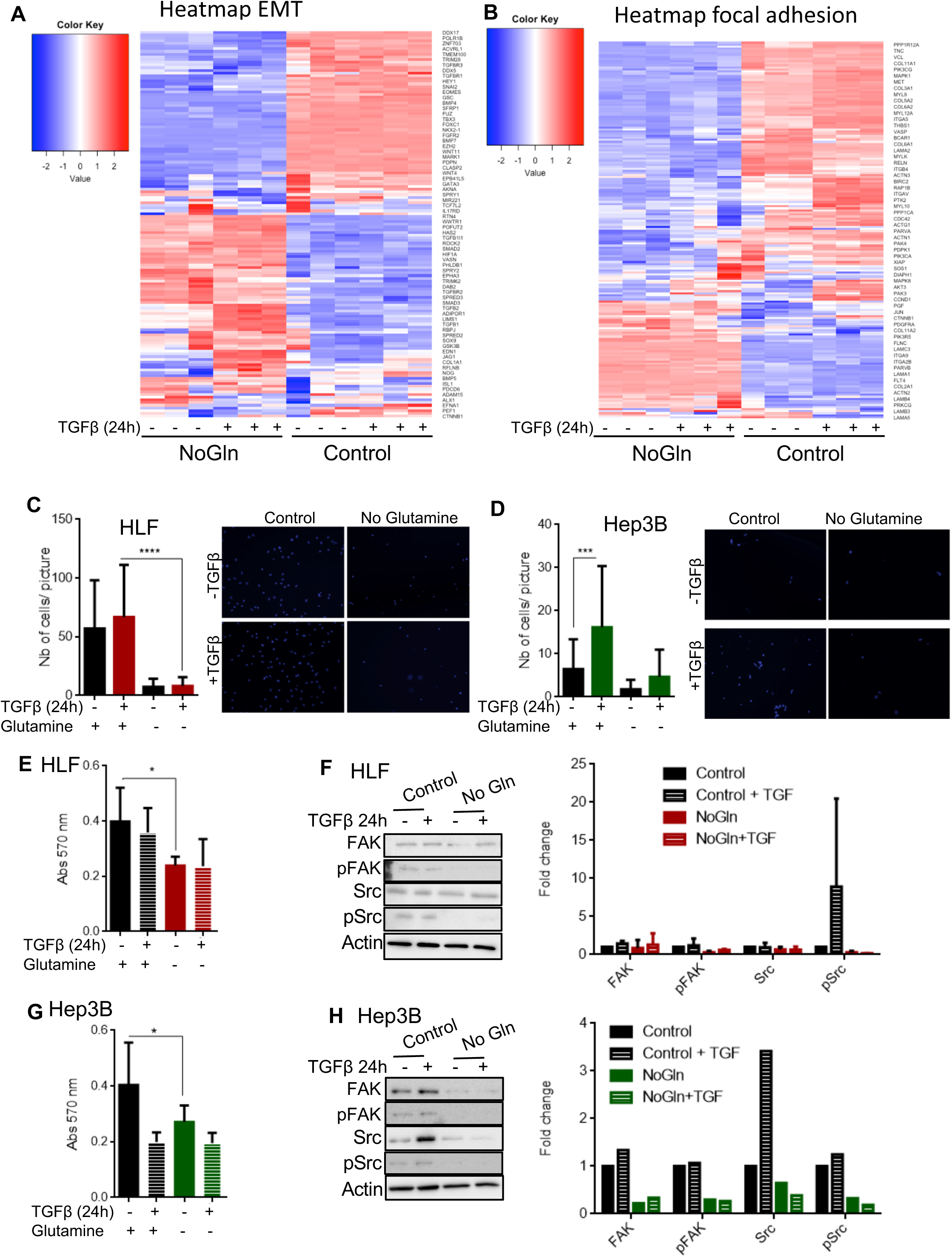
Glutamine deprivation alters cell invasion and adhesion in HCC. A,B. Heat-map of the Gene Ontology EMT related genes- and KEGG pathway focal adhesion-related genes in HLF cells under control or glutamine-deprived (no Gln) culture conditions with or without TGFβ treatment for 24 h. C, D. Transwell-based collagen invasion assays of HLF (C) and Hep3B cells (D) under control or glutamine-deprived (no Gln) culture conditions. Cells were treated or not with TGFβ for 15 h during the invasion period. E, F. Collagen-based adhesion assay of HLF (E) and Hep3B cells (F) under control or glutamine-deprived (no Gln) culture conditions. Cells were pretreated or not with TGFβ for 24 h before seeding. G, H. Immunoblot and quantification for the indicated proteins (FAK, pFAK, Src, pSrc, tubulin) in HLF (G) and Hep3B cells (H) under control or glutamine-deprived (no Gln) culture conditions with ot without TGFβ treatment for 24 h. Each graph represents data from (n=3 for HLF, n=1 for Hep3B) biological replicates as means with SEM and associated significance as: * p<0.05, ** p<0.01, *** p<0.001, **** p<0.0001.

## DISCUSSION

Tumors require glutamine to produce energy and thereby sustain cell survival and malignancy. In solid tumors, the access to glutamine is easier for the cells residing on the edge of the tumor, which also are the cells that detach from the tumors and invade surrounding tissues. Our study shows that glutamine is needed to maintain a certain proliferative rate, clonogenic and self-renewal capacities (Fig. 2). More interestingly, glutamine metabolism seems to be needed to maintain a mesenchymal phenotype with invasive capacities (Fig. 4, 5). Indeed, the absence of glutamine in the HCC culture medium led to a reinforcement of epithelial phenotypes (Hep3B) or a switch from mesenchymal to epithelial phenotype (HLF) associated with a dramatic decrease of mesenchymal gene expression (Fig. 5) and a loss of invasive capacity of the cells (Fig. 6). We therefore conclude that glutamine deprivation pushes the cells to become more epithelial, this effect being logically more obvious in mesenchymal cells.

TGFβ is a master signaling molecule that regulates the epithelial-to-mesenchymal transition, a process by which the cells migrate and invade the surrounding tissues by breaking their cell-cell junctions and by reorganizing their interaction with the extracellular matrix. RNA-sequencing data and gene ontology analysis in HLF treated with TGFβ followed by functional assays confirm that the activation of the TGFβ pathways (Fig. 5), the number of TGFβ-regulaed genes (Supplementary fig.3), as well as the expression of its target genes related to ECM structure and cell-cell adhesion (Fig. 4, 6) are reduced by the lack of glutamine.

It is known that glutamine metabolism affects cancer cell aggressiveness via its effect on stemness, migration, on the regulation of the redox state or its effect on cell proliferation. It is also known that glutamine metabolism activity is up-regulated in cancer by the up-regulation of key regulators of glutaminolysis such as glutamine transporters, glutaminase and lactate dehydrogenase A by TGFβ in HCC (Soukupova *et al*, 2017), or c-Myc in glioma and mouse embryonic fibrobalsts (Wise *et al*, 2008). However, the regulation of gene expression by long term glutamine starvation has not been previously established. The lack of nutrients triggers the autophagy as an intracellular source of energy for short-term survival. In normal conditions, the metabolic regulator mTORC1 is active and negatively regulates autophagy via the inhition of the autophagy-essential kinase Unc-51-like kinase-1 (ULK1) by phosphorylation (Altman *et al*, 2014). Under metabolic stress, for instance in nutrient-deprived condition, mTORC1 is inhibited and dissocates from ULK1, which is now able to phosphorylate and activate PI3KC complex that triggers autophagy. The inactivation of mTORC1 by the lack of nutrients is the main regulator of this process (Nobukuni *et al*, 2005). In the context of glutamine metabolism, a high mitochondrial activity maintained by a great intake of glutamate, produces a sufficient quantity of α-ketoglutarate to enter the TCA cycle and exit the mitochondria to maintain the mTORC1 activity. When the glutamine metabolism is reduced, α-ketoglutarate is only used to fuel the TCA cycle which leads to the inactivation of mTORC1 and therefore initiates autophagy (Durán *et al*, 2012). It is reported that TGFβ signaling activates mTORC1 via the activation of the PI3K-Akt pathway. The activation of mTORC1 by TGFβ triggers the metabolic reprogramming of lung fibroblasts to myoblasts via the activation of ATF4 (O’Leary *et al*, 2020) or impairs cell-cell adhesion and polarization in keratinocytes by mediating Rho kinase activation in epithelial cells (Asrani *et al*, 2017). One possible explanation to the alteration of TGFβ signaling by glutamine-deprivation would be that the lack of glutamine inactivates the mTORC1 in order to initiate autophagy. In this condition, it is possible that TGFβ can no longer activate mTORC1 which in turns leads to impaired TGFβ signaling.

## Additional Information

Additional information includes Additional files 1 and 2 (Tables S1, S2, S3, S4 and S5) and Additional file 3 (seven figures).

## Declarations

### Ethics approval and consent to participate

Not applicable.

### Consent for publication

Not applicable.

### Availability of data and materials

Further details or additional materials are available upon reasonable request.

## ACKNOWLEDGEMENTS

We thank other members of our groups for assistance and advice during the project. RNA-sequencing was performed by the SNP&SEQ Technology Platform at the National Genomics Infrastructure Sweden and Science for Life Laboratory. The SNP&SEQ Platform is also supported by the Swedish Research Council and the Knut and Alice Wallenberg Foundation.

This work was funded by: Ludwig Cancer Research, Swedish Cancer Society (CAN 2015/438, CAN2018/469, CAN2021/1506Pj01H to AM), Swedish Research Council (2017-01588, 2018-02757 to AM, 2015-02757, 2020-01291 to CHH), European Research Council (787472 to CHH), STINT (IB 2019-8148), OE och Elda Johanssons stiftelse (2020), Lars Hiertas (FO2020-033) and Magnus Bergvalls Stiftelse (2020-03781) to LC. The APC was funded by Uppsala University.

## Author contributions

L.C. conceived the project. C.G. and L.C. designed the experiments. C.G., S.C., I.C.G., T.N.B. and L.C. acquired the data. C.G., S.C., A.C., P.S. and L.C. analyzed the data. C.G., C-H.H., A.M., P.S., and L.C interpreted the data. C.G. drafted the article. A.M. and L.C. revised the article draft. All authors critically revised the article for important intellectual content and provided final approval prior to submission for publication.

## Abbreviations

ATP: adenosine triphosphate
CAF: cancer associated fibroblast
ECM: extracellular matrix
FAK: focal adhesion kinase
Gln: Glutamine
GLS: glutaminase
HCC: hepatocellular carcinoma
MMP: matrix metalloproteases
ROS: reactive oxygen species
TCA: tricarboxylic acid
TGF-β: transforming growth factor β

## REFERENCES

Albrecht J, Sidoryk-Wegrzynowicz M, Zielinska M, Aschner M (2010) Roles of glutamine in neurotransmission. Neuron Glia Biol 6: 263–276

Altman JK, Szilard A, Goussetis DJ, Sassano A, Colamonici M, Gounaris E, Frankfurt O, Giles FJ, Eklund EA, Beauchamp EM et al (2014) Autophagy is a survival mechanism of acute myelogenous leukemia precursors during dual mtorc2/mtorc1 targeting. Clin Cancer Res 20: 2400–2409

Asrani K, Sood A, Torres A, Georgess D, Phatak P, Kaur H, Dubin A, Talbot CC, Jr., Elhelu L, Ewald AJ et al (2017) Mtorc1 loss impairs epidermal adhesion via tgf-β/rho kinase activation. J Clin Invest 127: 4001–4017

Boroughs LK, DeBerardinis RJ (2015) Metabolic pathways promoting cancer cell survival and growth. Nat Cell Biol 17: 351–359

Bray F, Ferlay J, Soerjomataram I, Siegel RL, Torre LA, Jemal A (2018) Global cancer statistics 2018: Globocan estimates of incidence and mortality worldwide for 36 cancers in 185 countries. CA Cancer J Clin 68: 394–424

Caja L, Dituri F, Mancarella S, Caballero-Diaz D, Moustakas A, Giannelli G, Fabregat I (2018) Tgf-β and the tissue microenvironment: Relevance in fibrosis and cancer. Int J Mol Sci 19

Caruso S, Calatayud AL, Pilet J, La Bella T, Rekik S, Imbeaud S, Letouze E, Meunier L, Bayard Q, Rohr-Udilova N et al (2019) Analysis of liver cancer cell lines identifies agents with likely efficacy against hepatocellular carcinoma and markers of response. Gastroenterology 157: 760–776

Chen Y, Lun AT, Smyth GK (2016) From reads to genes to pathways: Differential expression analysis of rna-seq experiments using rsubread and the edger quasi-likelihood pipeline. F1000Res 5: 1438

Coulouarn C, Factor VM, Thorgeirsson SS (2008) Transforming growth factor-β gene expression signature in mouse hepatocytes predicts clinical outcome in human cancer. Hepatology 47: 2059–2067

Crosas-Molist E, Bertran E, Sancho P, Lopez-Luque J, Fernando J, Sanchez A, Fernandez M, Navarro E, Fabregat I (2014) The nadph oxidase nox4 inhibits hepatocyte proliferation and liver cancer progression. Free Radic Biol Med 69: 338–347

Dobin A, Davis CA, Schlesinger F, Drenkow J, Zaleski C, Jha S, Batut P, Chaisson M, Gingeras TR (2013) Star: Ultrafast universal rna-seq aligner. Bioinformatics 29: 15–21

Duran RV, Oppliger W, Robitaille AM, Heiserich L, Skendaj R, Gottlieb E, Hall MN (2012) Glutaminolysis activates rag-mtorc1 signaling. Molecular cell 47: 349–358

Durán RV, Oppliger W, Robitaille AM, Heiserich L, Skendaj R, Gottlieb E, Hall MN (2012) Glutaminolysis activates rag-mtorc1 signaling. Molecular cell 47: 349–358

Fan J, Kamphorst JJ, Mathew R, Chung MK, White E, Shlomi T, Rabinowitz JD (2013) Glutamine-driven oxidative phosphorylation is a major atp source in transformed mammalian cells in both normoxia and hypoxia. Mol Syst Biol 9: 712

Friedl P, Alexander S (2011) Cancer invasion and the microenvironment: Plasticity and reciprocity. Cell 147: 992–1009

Gameiro PA, Struhl K (2018) Nutrient deprivation elicits a transcriptional and translational inflammatory response coupled to decreased protein synthesis. Cell reports 24: 1415–1424

Garcia-Gomez P, Golán I, M SD, Mezheyeuski A, Bellomo C, Tzavlaki K, Morén A, Carreras-Puigvert J, Caja L (2022) Nox4 regulates tgfβ-induced proliferation and self-renewal in glioblastoma stem cells. Mol Oncol 16: 1891–1912

Hanahan D, Weinberg RA (2011) Hallmarks of cancer: The next generation. Cell 144: 646–674

Haussinger D, Schliess F (2007) Glutamine metabolism and signaling in the liver. Front Biosci 12: 371–391

Jin X, Zhang S, Wang N, Guan L, Shao C, Lin Y, Liu J, Li Y (2022) High expression of tgf-β1 contributes to hepatocellular carcinoma prognosis via regulating tumor immunity. Front Oncol 12: 861601

Ju HQ, Ying H, Tian T, Ling J, Fu J, Lu Y, Wu M, Yang L, Achreja A, Chen G et al (2017) Mutant kras- and p16-regulated nox4 activation overcomes metabolic checkpoints in development of pancreatic ductal adenocarcinoma. Nat Commun 8: 14437

Kamphorst JJ, Cross JR, Fan J, de Stanchina E, Mathew R, White EP, Thompson CB, Rabinowitz JD (2013) Hypoxic and ras-transformed cells support growth by scavenging unsaturated fatty acids from lysophospholipids. Proc Natl Acad Sci U S A 110: 8882–8887

Klepinin A, Zhang S, Klepinina L, Rebane-Klemm E, Terzic A, Kaambre T, Dzeja P (2020) Adenylate kinase and metabolic signaling in cancer cells. Front Oncol 10: 660

Lassmann T, Hayashizaki Y, Daub CO (2011) Samstat: Monitoring biases in next generation sequencing data. Bioinformatics 27: 130–131

Liao Y, Smyth GK, Shi W (2014) Featurecounts: An efficient general purpose program for assigning sequence reads to genomic features. Bioinformatics 30: 923–930

Mazat JP, Devin A, Ransac S (2020) Modelling mitochondrial ros production by the respiratory chain. Cell Mol Life Sci 77: 455–465

Mestre-Farrera A, Bruch-Oms M, Pena R, Rodriguez-Morato J, Alba-Castellon L, Comerma L, Quintela-Fandino M, Dunach M, Baulida J, Pozo OJ et al (2021) Glutamine-directed migration of cancer-activated fibroblasts facilitates epithelial tumor invasion. Cancer Res 81: 438–451

Moustakas A, Heldin C-H (2009) The regulation of tgfβ signal transduction. Development 136: 3699–3714

Nobukuni T, Joaquin M, Roccio M, Dann SG, Kim SY, Gulati P, Byfield MP, Backer JM, Natt F, Bos JL et al (2005) Amino acids mediate mtor/raptor signaling through activation of class 3 phosphatidylinositol 3oh-kinase. Proc Natl Acad Sci U S A 102: 14238–14243

O’Leary EM, Tian Y, Nigdelioglu R, Witt LJ, Cetin-Atalay R, Meliton AY, Woods PS, Kimmig LM, Sun KA, Gokalp GA et al (2020) Tgf-β promotes metabolic reprogramming in lung fibroblasts via mtorc1-dependent atf4 activation. Am J Respir Cell Mol Biol 63: 601–612

Penuelas-Haro I, Espinosa-Sotelo R, Crosas-Molist E, Herranz-Iturbide M, Caballero-Diaz D, Alay A, Sole X, Ramos E, Serrano T, Martinez-Chantar ML et al (2022) The nadph oxidase nox4 regulates redox and metabolic homeostasis preventing hcc progression. Hepatology

Prasad P, Roy SS (2021) Glutamine regulates ovarian cancer cell migration and invasion through ets1. Heliyon 7: e07064

Soukupova J, Malfettone A, Hyrossova P, Hernandez-Alvarez MI, Penuelas-Haro I, Bertran E, Junza A, Capellades J, Giannelli G, Yanes O et al (2017) Role of the transforming growth factor-β in regulating hepatocellular carcinoma oxidative metabolism. Scientific reports 7: 12486

Torrino S, Grasset EM, Audebert S, Belhadj I, Lacoux C, Haynes M, Pisano S, Abelanet S, Brau F, Chan SY et al (2021) Mechano-induced cell metabolism promotes microtubule glutamylation to force metastasis. Cell Metab 33: 1342–1357 e10

Tsubakihara Y, Moustakas A (2018) Epithelial-mesenchymal transition and metastasis under the control of transforming growth factor β. Int J Mol Sci 19

Tzavlaki K, Moustakas A (2020) Tgf-β signaling. Biomolecules 10

Warburg O (1956) On respiratory impairment in cancer cells. Science 124: 269–270

Wendt MK, Schiemann WP (2009) Therapeutic targeting of the focal adhesion complex prevents oncogenic tgf-β signaling and metastasis. Breast Cancer Res 11: R68

Wise DR, DeBerardinis RJ, Mancuso A, Sayed N, Zhang XY, Pfeiffer HK, Nissim I, Daikhin E, Yudkoff M, McMahon SB et al (2008) Myc regulates a transcriptional program that stimulates mitochondrial glutaminolysis and leads to glutamine addiction. Proc Natl Acad Sci U S A 105: 18782–18787

Yoo HC, Yu YC, Sung Y, Han JM (2020) Glutamine reliance in cell metabolism. Exp Mol Med 52: 1496–1516

Yu G, Wang LG, Han Y, He QY (2012) Clusterprofiler: An r package for comparing biological themes among gene clusters. OMICS 16: 284–287

