## Supplementary information for "Glutamine deprivation alters TGF-β signaling in hepatocellular carcinoma"

### SUPPLEMENTAL TABLES

**Supplementary Table S1.** List of antibodies used with dilution factors and application

| **Antibody** | **Source** | **Specification** |
|---|---|---|
| anti-Smad2 (1:1000 immunoblotting) | Cell Signaling Technology, Danvers, MA, USA | Cat# 5339S |
| anti-pSmad2 (1:500 immunoblotting) | Ludwig Cancer Research-Uppsala Branch, Sweden | Home-made |
| anti-ERK (1:1000 immunoblotting) | Cell Signaling Technology, Danvers, MA, USA | Cat# 4695 |
| anti-pERK (1:1000 immunoblotting) | Cell Signaling Technology, Danvers, MA, USA | Cat# 4370S |
| anti-Akt (1:1000 immunoblotting) | Cell Signaling Technology, Danvers, MA, USA | Cat# C67E7 |
| anti-pAkt (1:1000 immunoblotting) | Cell Signaling Technology, Danvers, MA, USA | Cat# C31E5E |
| anti-ECadherin (1:1000 immunoblotting, 1:200 immunofluorescence) | Cell Signaling Technology, Danvers, MA, USA | Cat# 24E10 |
| anti-Fibronectin (1:1000 immunoblotting, 1:200 immunofluorescence) | Sigma-Aldrich AB, Stockholm, Sweden | Cat# F3648 |
| anti-Vimentin (1:1000 immunoblotting, 1:200 immunofluorescence) | Cell Signaling Technology, Danvers, MA, USA | Cat# 5741S |
| anti-PAI1 | Abcam, Cambridge, UK | Cat# ab66705 |

| | | |
|---|---|---|
| (1:1000 immunoblotting) | | |
| anti-Tubulin (1:1000 immunoblotting) | Santa Cruz | Cat# sc-8035 |
| anti-SRC (36D10) (1:1 000 immunoblotting) | Cell signaling Technology, Dancers, MA, USA | Cat# 2109 |
| anti-SRC pY419 (1:1000 immunoblotting) | Cell signaling Technology, Dancers, MA, USA | Cat# 2101 |
| anti-FAK (1:1000 immunoblotting) | BD Biosciences | Cat# 610087 |
| anti-FAK pY397 (1:1000 immunoblotting) | BD Biosciences | Cat# 611722 |
| anti-CyclinE (1:1000 immunobotting) | Cell signaling Technology, Dancers, MA, USA | Cat# 4129T |
| anti-PARP1 (1:1000 immunoblottig) | Cell signaling Technology, Danvers, MA, USA | Cat# 5625 |
| anti-Caspase3 (1:1000 immunobloting) | Cell signaling Technology, Dancers, MA, USA | Cat# 9662 |
| anti-HP95 (1:2000 immunoblotting) | Sigma-Aldrich AB | Cat# HPA011905 |
| anti-Vinculin (AC-15) (1:200 immunofluorescence) | Sigma-Aldrich AB | Cat# V9131 |
| anti-Rabbit IgG H+L) Secondary, HRP | Thermo Fisher Scientific | Cat# 65-6120 |
| anti-Mouse IgG (H+L) Secondary, HRP | Thermo Fisher Scientific | Cat# 62-6520 |

**Supplementary Table S2.** List of oligonucleotides (Fw, forward; Rev, reverse)

| **mRNA-specific RT-PCR primers** | **Sequence** |
|---|---|
| *GLS1* | Fw GAAAGAGTACTGAGCCCTGAAG<br>Rev GGACAACTAAAAGAATGCCCC |
| *GLS2* | Fw TCCACAACTATGACAACCTGAG<br>Rev GCTGAGACATCGCCACTATAG |
| *SLC1A5* | Fw CCCTCATCTACTTCCTCTTCAC<br>Rev TTATTCTCCTCCACGCACTTC |
| *SLC7A5* | Fw TCTTCAACTGGCTCTGCG<br>Rev GAAGGAGACGGCGATCAG |
| *Nox4* | Fw GCAGGAGAACCAGGAGATTG<br>Rev CACTGAGAAGTTGAGGGCATT |
| *Smad2* | Fw TGCCTTCGGTATTCTGCTCCCCA<br>Rev TGGCTGGCACCCTGCAACAG |
| *Smad3* | Fw GCAATATTCCAGAGACCCCACC<br>Rev TAGGTTTGGAGAACCTGCGTCC |
| *Smad4* | Fw TGAAGGACTGTTGCAGATAGCA<br>Rev TCCAGGTGGTAGTGCTGTTATG |
| *E-Cadherin* | Fw TACGCCTGGGACTCCACCTA<br>Rev CCAGAAACGGAGGCCTGAT |
| *N-Cadherin* | Fw TCATGCACATCCTTCGATAAGACT<br>Rev CCTGCTTCAGGCGTCTGTAGA |
| *Snail* | Fw CACTATGCCGCGCTCTTTC<br>Rev GCTGGAAGGTAAACTCTGGATTAGA |
| *Fibronectin* | Fw CATCGAGCGGATCTGGCCC<br>Rev GCAGCTGACTCCGTTGCCCA |
| *HPRT1* | Fw CCCTGGCGTCGTGATTAGT<br>Rev CACCCTTTCCAAATCCTCAGC |

**Supplementary Table S4.** Excel files listing the unprocessed data of the RNAseq analysis in control or glutamine-deprived HLF cells (treated or not treated with TGFβ for 24h).

### SUPPLEMENTARY FIGURE TITLES AND LEGENDS

**Supplementary figure 1 related to figure 2. Glutamine deprivation does not affect senescence and apoptosis but decreases cell cycle activity**

A, B. Immunoblots for the indicated proteins (cyclinE, PARP, total and cleaved caspase3, β-actin) in HLF (A) and Hep3B (B) cells under control or glutamine-deprived (no Gln) culture conditions. C, D. Phase-contrast images showing the β-galactosidase based senescence assay in HLF (C) and Hep3B (D) cells under control or glutamine-deprived (no Gln) culture conditions. Magnification: ×10.

**Supplementary figure 2 related to figure 2. Glutamine deprivation reduces the self-renewal capacity of HLF, SNU-499 and Hep3B cells**

A-C. Phase-contrast images showing the spheres obtained on low-attachment plate in HLF (A), SNU-499 (B) and Hep3B (C) cells under control or glutamine-deprived (no Gln) culture conditions. Magnification bars (100 μm).

**Supplementary figure 3 related to figure 4. Transcriptomic analysis of glutamine-deprived HLF cells and after stimulation with TGFβ**

A. Volcano plots of differentially-regulated genes in HLF cells, in response to glutamine deprivation. Log fold-change threshold was set to +1 and -1; significantly regulated genes are shown in green, not significantly regulated genes are shown in red. B.Volcano plots of differentially-regulated genes, in response to glutamine deprivation in HLF cells and after TGFβ treatment for 24 h. Log fold-change threshold was set to +1 and -1; significantly regulated genes are shown in green, not significantly regulated genes are shown in red.  C. Volcano plots of differentially-regulated genes in HLF cells after TGFβ treatment for 24 h. Fold-change threshold was set to +2 and -2; significantly regulated genes are shown in green, not significantly regulated genes are shown in red. D. Volcano plots of differentially-regulated genes in glutamine-deprived HLF cells after TGFβ treatment for 24 h. Fold-change threshold was set to +2 and -2; significantly regulated genes are shown in green, not significantly regulated genes are shown in red.

**Supplementary figure 4 related to figure 6. Glutamine deprivation affects the expression of EMT-related proteins regulated by TGFβ**

A-D. Representative immunofluorescence staining pictures of fibronectin, E-cadherin, vimentin and vinculin (red) in HLF cells. Nuclei are stained in blue with DAPI. Magnification bars (50 μm).

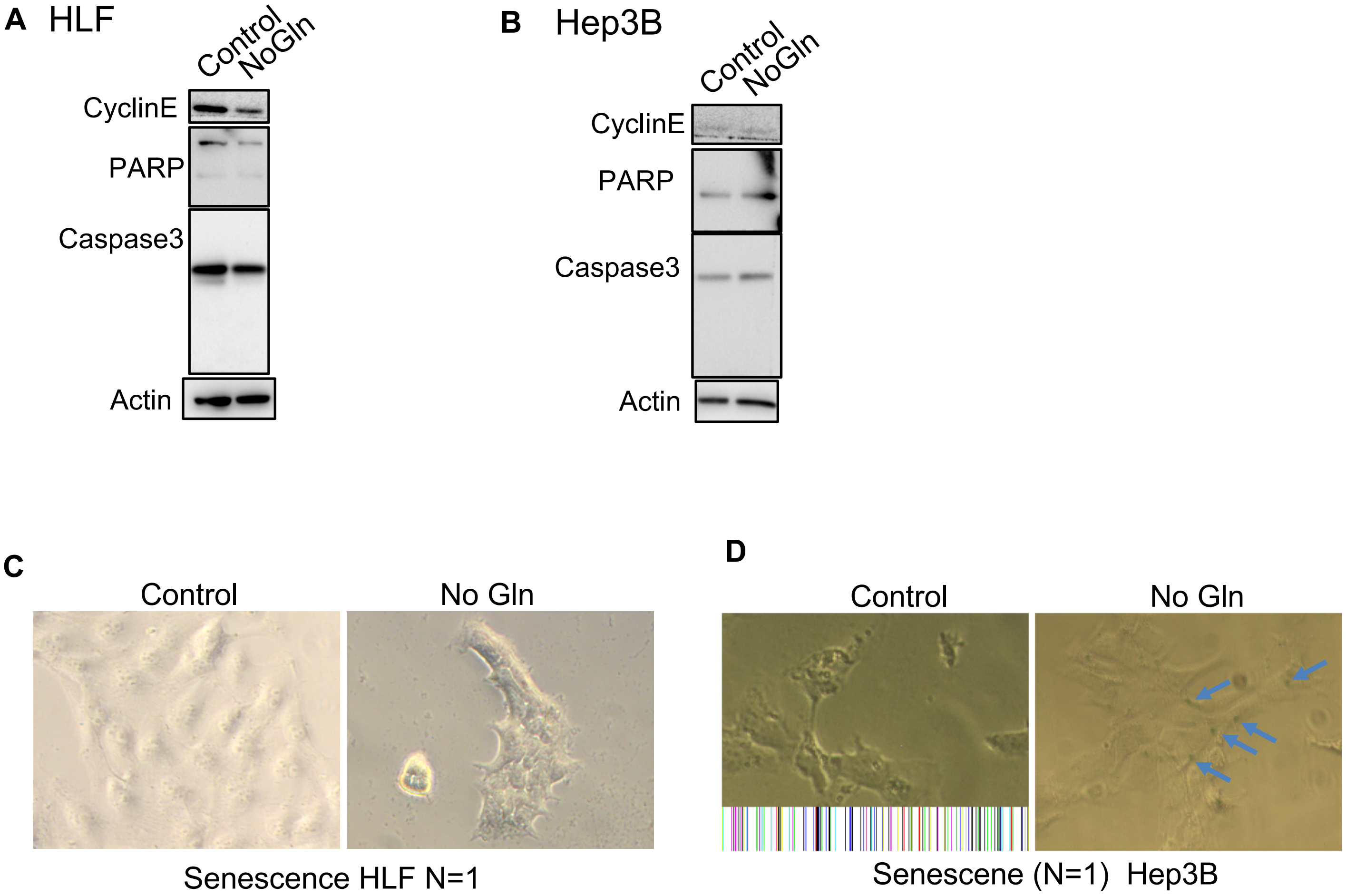


Supplementary figure 1 related to Figure 2

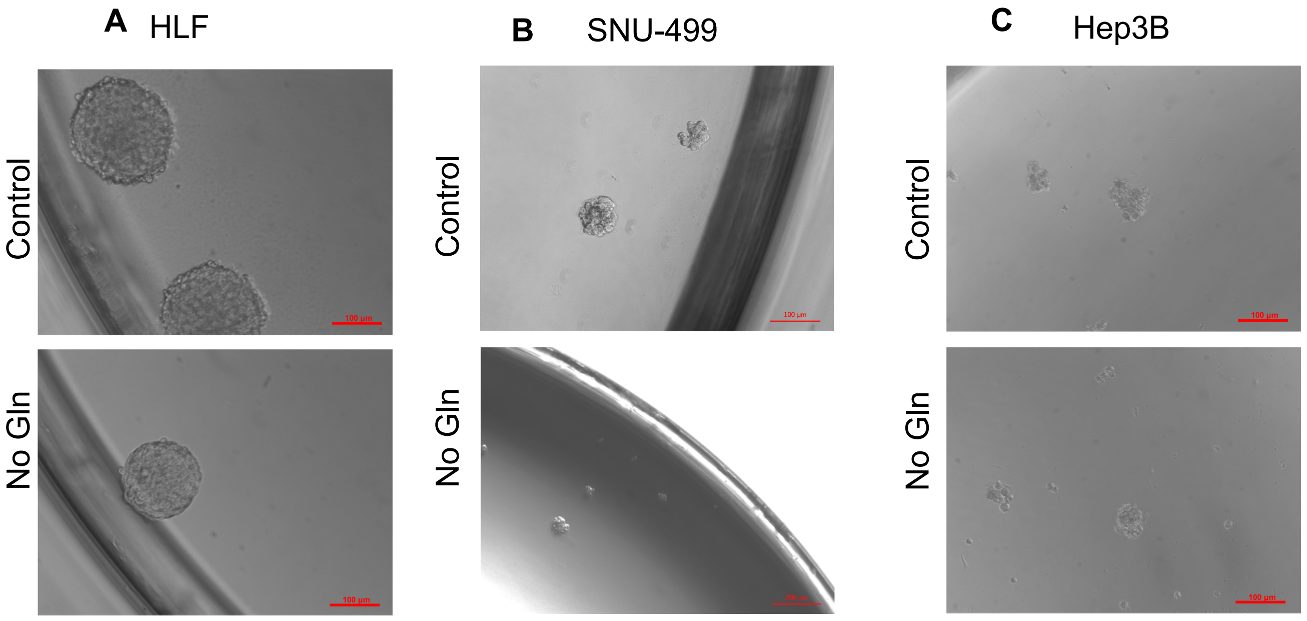


Supplementary figure 2 related to Figure 2

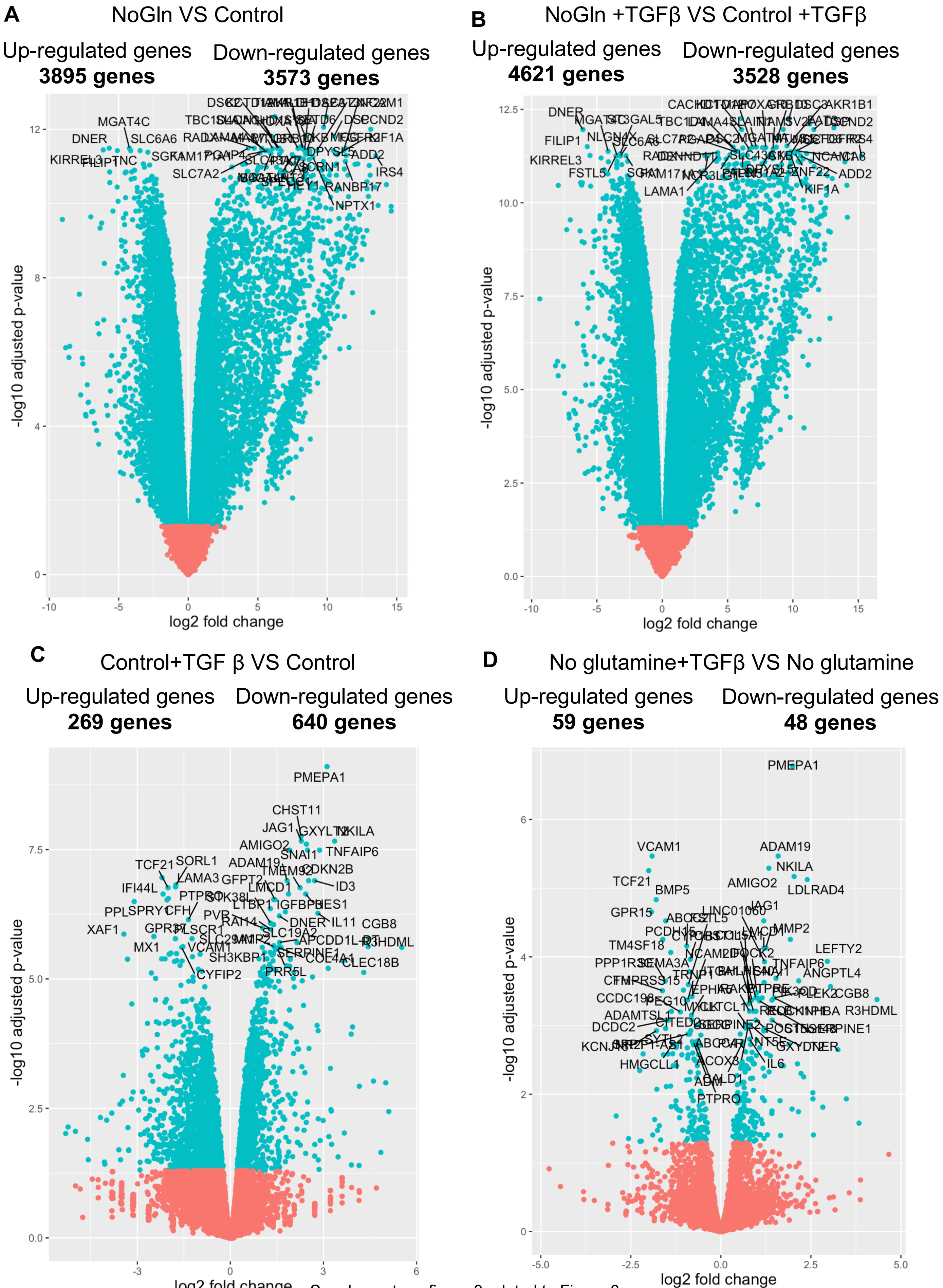


Supplementary figure 3 related to Figure 3

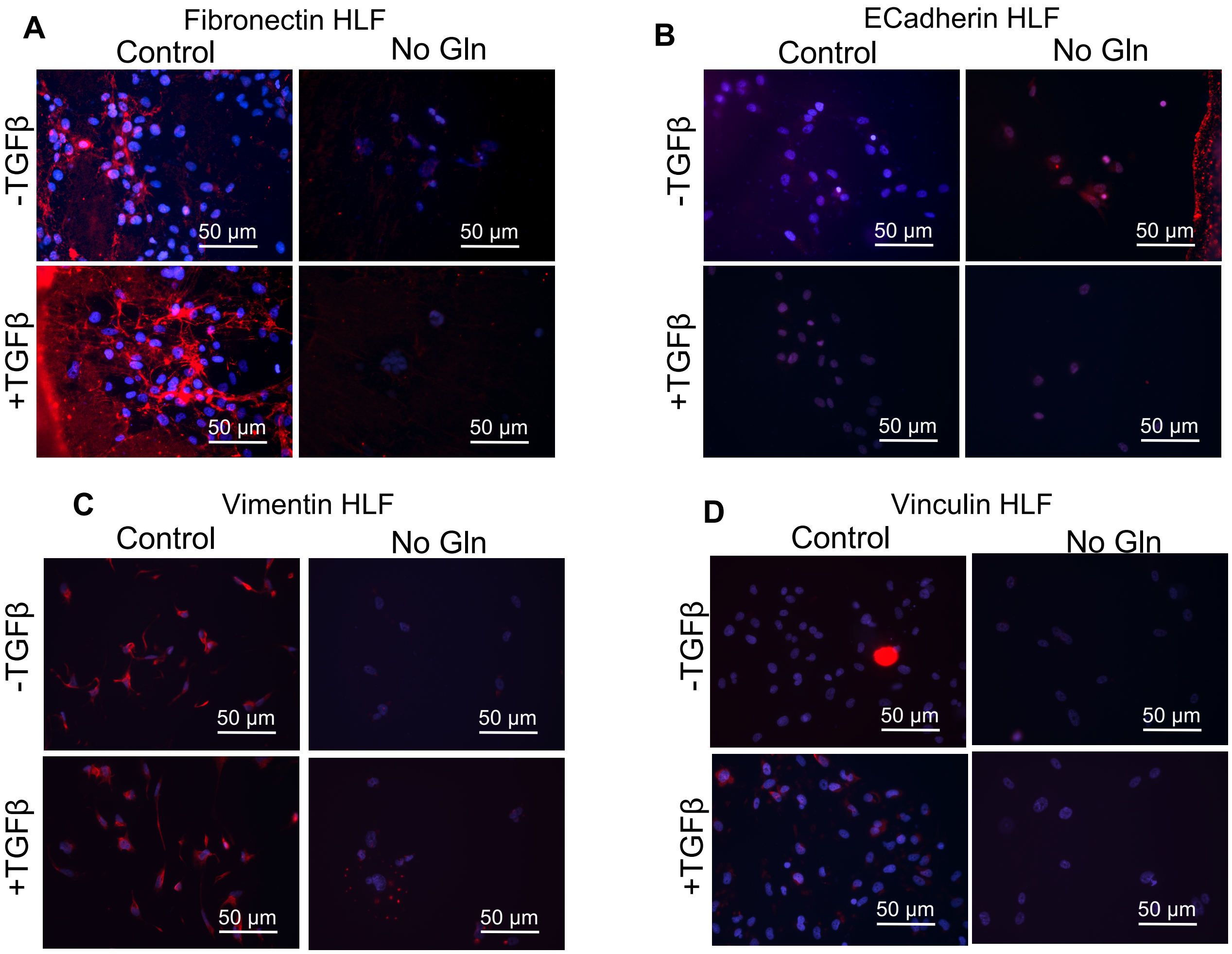


Supplementary figure 4 related to Figure 5
